# Disrupted Maternal Behavior in Morphine-Dependent Pregnant Rats and Anhedonia in their Offspring

**DOI:** 10.1101/2024.12.30.630830

**Authors:** Christopher T. Searles, Meghan E. Vogt, Iyanuoluwa Adedokun, Anne Z. Murphy

## Abstract

It is currently estimated that every 15 minutes an infant is born with opioid use disorder and undergoes intense early life trauma due to opioid withdrawal. Clinical research on the long-term consequences of gestational opioid exposure reports increased rates of social, conduct, and emotional disorders in these children. Here, we investigate the impact of perinatal opioid exposure (POE) on behaviors associated with anhedonia and stress in male and female Sprague Dawley rats. Young adult female rats were administered morphine via programmable, subcutaneous micro-infusion pumps before, during, and through one week post gestation. Maternal behavior was examined for fragmentation and entropy for the first two postnatal weeks; offspring were assessed for sucrose preference, social behavior, and stress responsivity. Overall, dams that received morphine across gestation displayed significantly less pup-directed behavior with increased fragmentation for nursing and higher entropy scores. In adolescence, male and female rat offspring exposed to morphine displayed reduced sucrose preference and, as adults, spent significantly less time socially interacting with familiar conspecifics. Changes in social behaviors were linked to increased activity in nondopaminergic mesolimbic reward brain regions. Although no treatment effects were observed in forced swim test performance, corticosterone levels were significantly increased in POE adult males. Together, these results suggest that perinatal morphine exposure results in anhedonic behavior, possibly due to fragmented and unpredictable maternal behavior in opioid-dependent dams.

**Significance Statement:** Clinical and preclinical research has shown that early life stress can produce lifelong depression-like symptoms. Here, we show that maternal opioid use during pregnancy produces depression-like symptoms in the offspring. We also observed disrupted maternal behavior in morphine-exposed dams that is similar to the changes in maternal behavior reported following other forms of early life stress. We propose that the stress of opioid exposure and suboptimal rearing leads to reduced sucrose preference, increased stress reactivity, and social avoidance behavior in the offspring, all of which are characteristics of anhedonia.

## Introduction

Across multiple decades, the United States has generally outpaced other nations in opioid use and related deaths, leading to a crisis that has only recently garnered the clinical attention it deserves (Pierce et al., 2021). One of the most significant at-risk groups for deleterious opioid consequences is pregnant women and their unborn children. Seven percent of women report using an opioid prescription during pregnancy, and of those women, one in five report misuse (Ko et al., 2020). As a consequence, between 2010 and 2017, the incidence rate of neonatal opioid withdrawal syndrome (NOWS) escalated across all races, income brackets, and insurance statuses from 4.0 to 7.3 in 1000 births (Hirai et al., 2021). While the immediate needs of these infants are well known and addressed in the hospital following delivery, the long-term behavioral and cognitive changes due to opioid exposure during pregnancy are understudied.

To address this gap in the literature, our lab has developed a novel perinatal opioid exposure (POE) model that administers daily pulsatile morphine to rat dams from before breeding to one week following parturition. Using this model, we have previously reported that gestational exposure to morphine reduces juvenile social play and alters both alcohol and sucrose intake during adolescence and early adulthood (Harder et al., 2023; Searles et al., 2023). Parallel changes in oxytocin and μ-opioid receptor expression in brain regions key for social behavior and reward signaling were also observed, suggesting an overall shift in reward processing. Similar results have been reported following early life manipulations in other models. For example, Molet et al. (2016) showed that housing rat dams and pups in impoverished environments for one week postnatally induced aberrant maternal behavior and anhedonia in male offspring. More specifically, male offspring that experienced early life stress had reduced sucrose preference and adolescent social play (Molet et al., 2016).

As opioid use during pregnancy is considered an “early life stressor” and our POE rats displayed aspects of anhedonia in their reduced social play and operant response for sucrose, we predicted our POE rats would show alterations in other hallmarks of anhedonia, namely, sucrose preference, adult social interaction, and forced swim. Additionally, due to the reported interaction between maternal care deficits in early life stress models and changes in reward-seeking behaviors and anhedonia in offspring, we also expanded upon our previously published analysis of maternal behavior to determine if POE induces suboptimal maternal care in more nuanced ways than simply gross time spent performing pup-directed actions (Bolton et al., 2018; Molet et al., 2016). Maternal care observations were used to calculate fragmentation, defined as a reduction in maternal care bout length, and entropy, or unpredictability in maternal care.

Based on the rationale above, the present study tested the hypothesis that perinatal morphine exposure leads to suboptimal maternal care, acting as an early life stressor and driving anhedonia in offspring.

## Materials and Methods

### Subjects

Female Sprague Dawley rats (age P60; Charles River Laboratories, Boston, MA) were used to generate male and female offspring. POE and control offspring were weaned at P21 into Optirat GenII individually ventilated cages (Animal Care Systems, Centennial, Colorado, USA) with corncob bedding in same-sex pairs or groups of three on a 12:12 hour light/dark cycle (lights on at 8:00 AM). Food (Lab Diet 5001 or Lab Diet 5015 for breeding pairs, St. Louis, MO, USA) and water were provided *ad libitum* throughout the experiment, except during maternal behavior recording and operant procedures. All studies were approved by the Institutional Animal Care and Use Committee at Georgia State University and performed in compliance with the National Institutes of Health Guide for the Care and Use of Laboratory Animals. All efforts were made to reduce the number of rats used in these studies and minimize pain and suffering.

### Perinatal opioid exposure design

Female Sprague Dawley rats were implanted with iPrecio SMP-200 microinfusion minipumps at postnatal day 60 (P60). Pumps were programmed to begin morphine or saline administration following one week of recovery from surgery. One week after morphine initiation, females were paired with sexually-experienced males for two weeks to induce pregnancy. Morphine (or saline) exposure continued throughout gestation. Rats were initially administered 10 mg/kg across three doses per day, with doses increasing weekly by 2 mg/kg/day until 16 mg/kg/day was reached. At approximately E18, pumps switched to twice-a-day dosing, as initial pilots using three-times-a-day dosing increased pup mortality early in life, possibly due to respiratory depression. Dams continued to receive decreasing doses of morphine up to P7. This protocol closely mirrors the clinical profile of a late adolescent female who develops an opioid use disorder, becomes pregnant, and continues using opioids throughout pregnancy. An essential part of this model is the administration of morphine prior to E15, the approximate date of μ-opioid receptor development in the fetal brain (Coyle & Pert, 1976). Pumps filled with sterile saline were used to generate control (CON) rats to account for surgery and pump refilling stress. See Figure 1 for the overall dosing and testing schedule.

**Figure 1.**
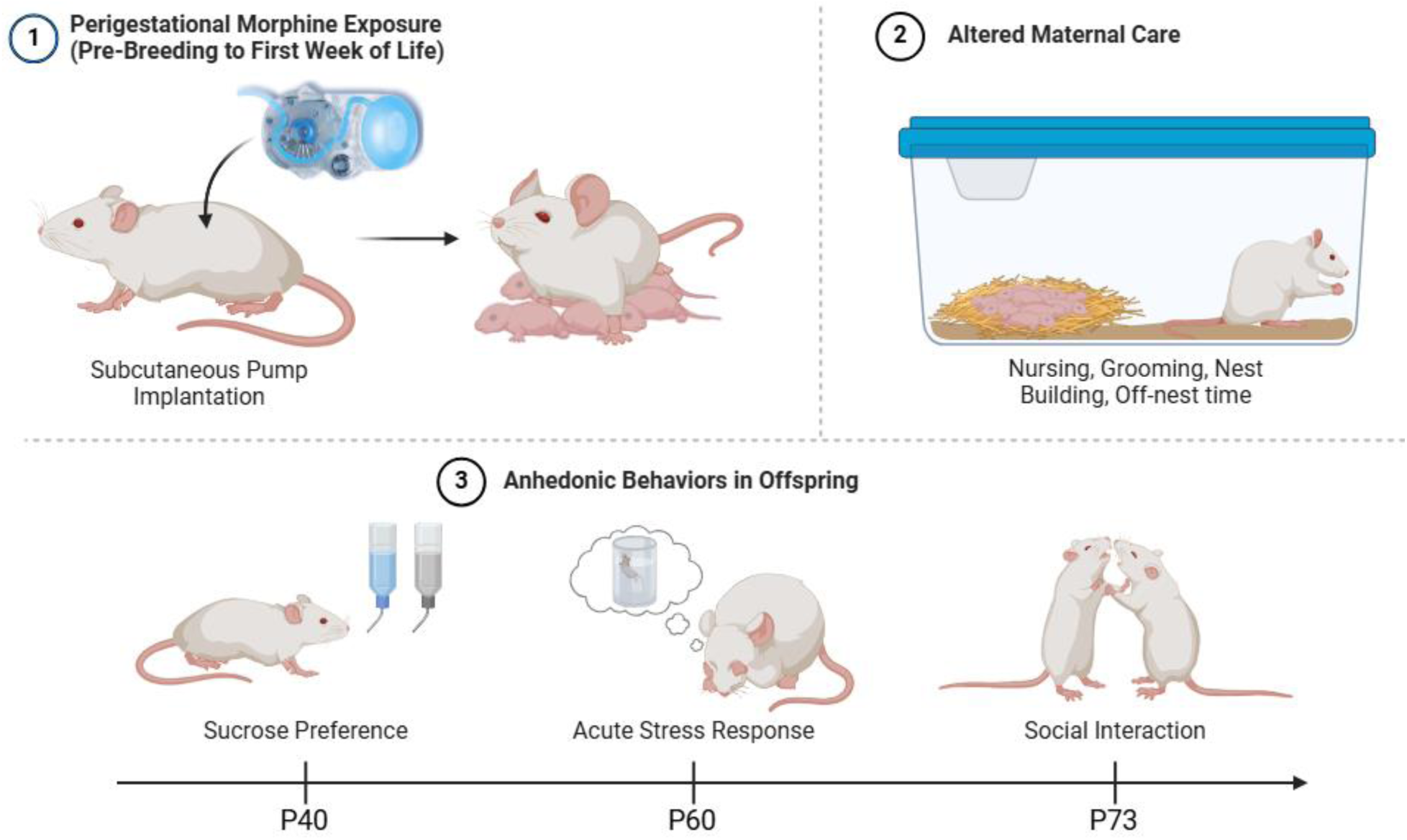
At P60, female Sprague Dawley rats are implanted with subcutaneous pumps to deliver morphine (or saline) prior to breeding through postnatal day 7 (P7). Maternal behavior was monitored at P2, P4, and P6 twice daily (before and after a scheduled infusion of morphine). Following morphine termination at P7, maternal behavior was observed once a day on P8, P10, P12, and P14. In adolescence, POE and CON rats were assessed for anhedonic behavior using sucrose preference test. Adult POE and CON rats were assessed for stress reactivity using FST and social behavior using social interaction test. Created with Biorender.

### Maternal behavior analysis

Rat maternal behaviors were observed on P2, P4, and P6 twice daily for 1hr: 2hr after lights on (prior to morphine infusion) and 6hr after lights on (after morphine infusion).

Following morphine cessation at P7, maternal behavior was observed once daily on P8, P10, P12, and P14, 4hr after lights on. All sessions were recorded for offline analysis. The following measures were scored: nursing, licking/grooming pups, nest building, self-grooming, returning pups to the nest, and time spent off nest. Time spent on individual behaviors was recorded and used to calculate behavior bout length to determine levels of fragmentation. A shorter average bout length for a type of behavior indicates higher fragmentation. Pattern analysis of behaviors was performed to calculate an entropy value, representing the predictability of behaviors. High predictability, where one behavior expectedly follows another, would be represented by a low entropy value. Calculation of entropy is described in depth by Molet et al. (2016).

### Sucrose preference test

At P40, POE and control rats were tested for sucrose preference using a 1% sucrose solution during their dark phase. On day 1, rats were individually housed, and their water bottles were replaced with two 200mL bottles of 1% sucrose for 24hr. On day 2, their water was returned for 6hr and then replaced with one water bottle and one sucrose bottle for 18hr. On day 3, rats were water-deprived for 12hr and then provided one water bottle and one sucrose bottle for 12hr. The positions of the bottles were counterbalanced. After 12hr of two-bottle-choice, the experimental water/sucrose bottles were removed, and rats were returned to their original home cages. Each day, all bottles were weighed pre– and post-ingestion to determine volume (mL) consumed.

### Forced swim test

Adult (P60) POE and control rats were tested for stress reactivity using the forced swim test (FST) (Victoria et al., 2014). An 80×200mm cylinder was filled with 25°C water deep enough that rats could touch the bottom while at the surface. All rats were given a 5min pre-swim one day prior to FST. On testing day, rats were placed in the water for a 5min FST; time immobile and latency to immobility were determined using ANY-Maze™ behavioral tracking software (Stoelting Co., Wood Dale, IL). Blood samples were collected from the saphenous vein into EDTA-hematology tubes (BD from Fisher Scientific) at four intervals relative to the FST to measure corticosterone levels: baseline (–60min), stress (5min), peak (30min), and recovery (75min). Blood samples were centrifuged at 3000 rcf at 4°C for 15 min, and plasma corticosterone was measured with ELISA (DetectX Corticosterone Multi-Format Kit, K014-H; Arbor Assays, Ann Arbor, MI). Correlation coefficients were measured against known standards (r^2^ > 0.98). Inter– and intra-assay CVs were below 12%.

### Social interaction test

At P73, POE and control rats were isolated for 24hr, then reunited with their cage mates in their original home cage. Behavior was recorded for 10min for offline analysis; time spent interacting, defined as sniffing, grooming, chasing, or playing, was measured.

### Immunohistochemistry

Ninety minutes following the social interaction test, one cohort of rats was sacrificed to assess forebrain and midbrain Fos and tyrosine hydroxylase (TH) expression. Rats were transcardially perfused with 0.9% sodium chloride saline containing 2% sodium nitrate followed by 4% paraformaldehyde fixative. Brains were extracted and stored in 4% paraformaldehyde at 4°C overnight, then transferred into 30% sucrose until sectioning. Fixed tissue was sectioned coronally at 25-µm with a Leica SM2010R microtome and stored at –20°C in cryoprotectant-antifreeze solution (Watson et al., 1986). Free-floating sections were rinsed in potassium phosphate-buffered saline (KPBS), incubated in 3% hydrogen peroxide for 30 minutes, and rinsed again in KPBS at room temperature. Tissue was then incubated in mouse anti-c-Fos primary antibody (Abcam Inc., Cambridge, MA, USA.; 1:10,000) overnight at room temperature. Tissue was then rinsed and incubated in biotinylated donkey anti-mouse IgG (Jackson Immunoresearch, West Grove, PA, USA; 1:600) for 1 hour. Secondary antibody was amplified with avidin-biotin solution for 1 hour. After tissue was rinsed with KPBS and sodium acetate (0.175M; pH 6.5), immunoreactivity was visualized using nickel sulfate 3,3′-diaminobenzidine solution (2 mg/10 ml) and 0.08% hydrogen peroxide in sodium acetate buffer. After 20 minutes, tissue was rinsed in sodium acetate and KPBS. Tissue was then blocked with 5% normal donkey serum (VWR, Inc., Pittsburgh, Pennsylvania, USA) for 1 hour at room temperature and incubated in sheep anti-TH primary antibody (Fisher Scientific, Inc., Palatine, IL, USA; 1:1000) for 1 hour at room temperature followed by 24 hours at 4°C. Primary antibody was rinsed using KPBS and incubated in donkey anti-sheep AlexaFluor 555 secondary antibody (Fisher Scientific, Palatine, IL, USA; 1:200) for 1.5 hours at room temperature. Tissue was rinsed and mounted on gelatin-subbed slides, air-dried, and cover-slipped with ProLong Diamond Antifade Mountant (Fisher Scientific, Inc., Palatine, IL, USA).

16-bit images were captured using a Keyence BZ-X700 microscope and software for immunohistochemistry quantification. Subregions of the medial prefrontal cortex (mPFC), nucleus accumbens (NAc), habenula, amygdala, and ventral tegmental area (VTA) were bilaterally imaged at 10x magnification (2-3 sections per rat, average of 4-6 values). Fos+ cells were detected automatically using ImageJ software and then manually validated. Total number of Fos+ cells was averaged to produce a mean number per rat per region. In VTA sections, TH immunoreactivity was used to determine if Fos+ cells were dopaminergic.

### Statistical analysis

All data met the requirements for parametric analyses (normality, homogeneity of variance). The effects of sex, treatment (morphine/saline), and session (where applicable) were assessed using two-or three-way mixed model ANOVA with an alpha level of 0.05. Given that repeated measures ANOVA cannot handle missing values, mixed models using Greenhouse-Geisser were implemented where outlier correction was necessary. Outliers were identified using GraphPad Prism’s ROUTS method. Post hoc testing was performed using Tukey’s or Sidak’s. Experiments involving repeated measures were not tested for daily individual group differences. No significant litter effects were observed; therefore, group data comprised multiple litters (6 control/7 POE for maternal behavior; 2 control/2 POE for offspring behaviors). GraphPad Prism 9.5.1 was used for all statistical analyses.

## Results

### Maternal Care

To determine if opioid use during pregnancy produced suboptimal maternal care, we first determined whether maternal care fragmentation differed between POE and CON dams as reflected by mean bout length of each behavior. Our analysis identified significant interactions of treatment and session for nursing bout length (F_treatment*session_(9,83)=2.53, p=0.01; Figure 2a). Post-hoc analysis showed that the mean bout length for nursing was significantly shorter in POE dams compared to CON at P2 PM and P4 PM assessments when circulating levels of morphine were high. To assess for patterns when morphine was present or absent, we also grouped data by category: low levels of morphine (P2-P6 AM), high levels of morphine (P2-P6 PM), and no morphine (P8-P14). Overall, the average nursing bout length was shorter for POE than CON during morphine highs (PM) (F_treatment*session_(2,21)=5.57, p=0.01; Figure 2b). No differences were noted during low morphine (AM) or no morphine (P8-P14). Interestingly, CON rat mothers displayed significantly longer nursing bouts in the afternoon (when POE morphine levels were high) compared to P8-P14 (no morphine). Although mean licking/grooming bout length was longer in POE dams across the majority of timepoints examined, these differences were not significant (Figure 2c). Similarly, no significant differences were noted in bout length when data were grouped by morphine level (Figure 2d). Bouts of self-grooming behavior were significantly different in POE rats compared to CON (F_treatment*session_(9,88)=2.03, p=0.04; Figure 2e). No significant differences in self-grooming behavior were noted when data were grouped by morphine level (Figure 2f), although bout length for POE dams was consistently longer than CON.

**Figure 2.**
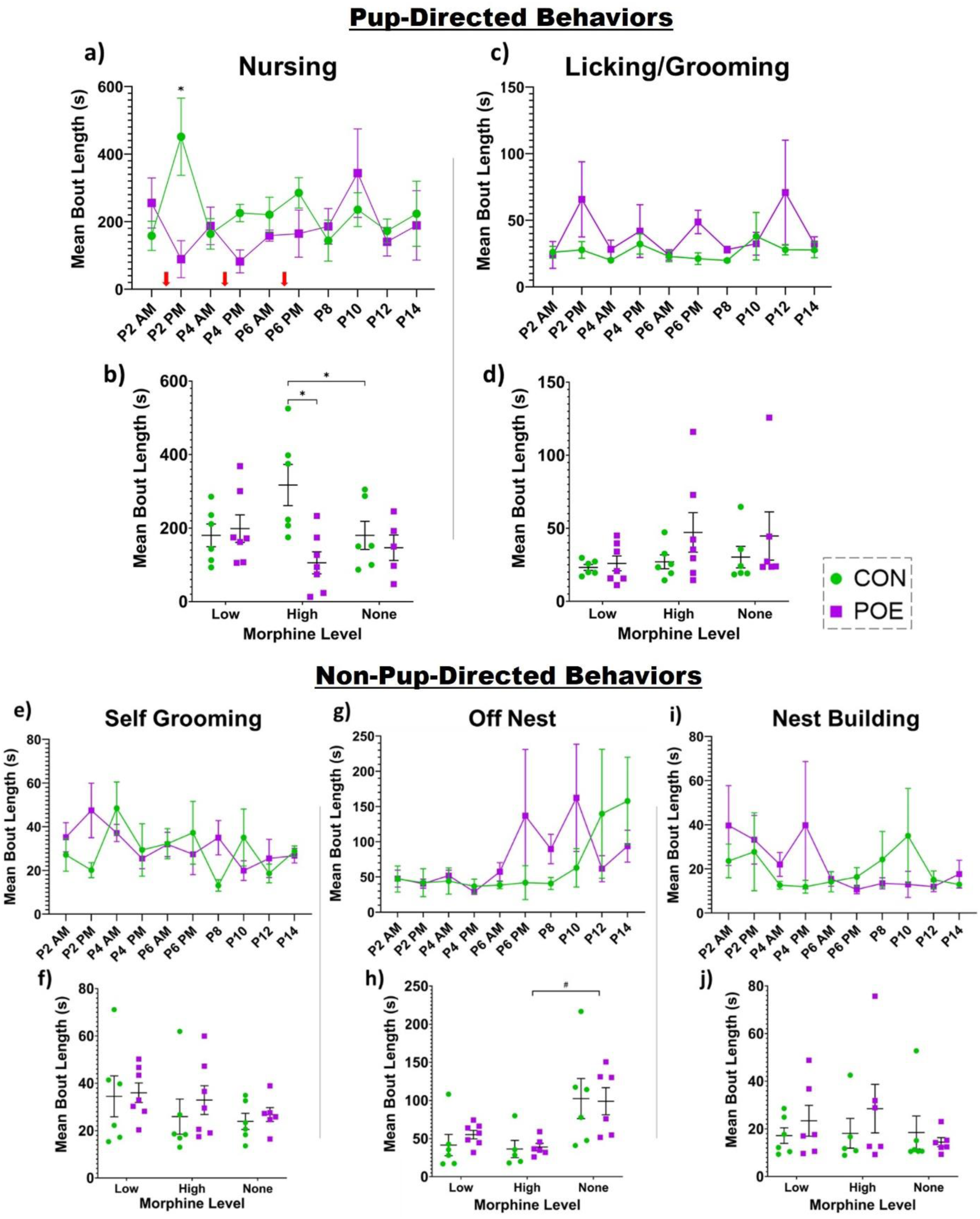
Maternal behavior was assessed for differences in bout length and unpredictability. Specifically, we measured nursing. (**a-b**), licking/grooming of pups (**c-d**), self-grooming (**e-f**), off-nest time (**g-h**), and nest building (**i-j**). Red arrows indicate when dams received a bolus of morphine. *Significant difference: Nursing (CON vs POE P2 PM and P4 PM; CON vs POE high morphine level; CON high vs none), Licking/Grooming (CON vs POE P6 PM and P8), Self-Grooming (CON vs POE P8); **#**Statistical trend (p<0.08): Off-Nest (CON vs POE P8; POE high vs none). Graphs indicate mean ± SEM.

There was no significant treatment effect for off-nest mean bout length (Figure 2g-h). Following morphine cessation at P8, POE dams displayed a trending increase (p=.07) in their off-nest bout length compared to CON. However, no significant interaction of treatment and session was observed (Figure 2g). No treatment effects were observed in nest building mean bout length (Figure 2i-j).

We also assessed total time spent performing each behavior across the three morphine levels. There were significant treatment and session effects on total time spent nursing as well as a significant session x treatment interaction, with all dams nursing the most during morning sessions and POE dams nursing less than CON dams, especially in the morning when morphine levels were low (F_session x treatment_(2,22)=6.919, p=0.0047; F_treatment_(1,11)=5.388, p=0.0405; F_session_(2,22)=159.4, p<0.0001; Figure 3a). There was a significant effect of session but no treatment effect on licking/grooming pups (F_session_(1.160,12.76)=36.62, p<0.0001; Figure 3b).

**Figure 3.**
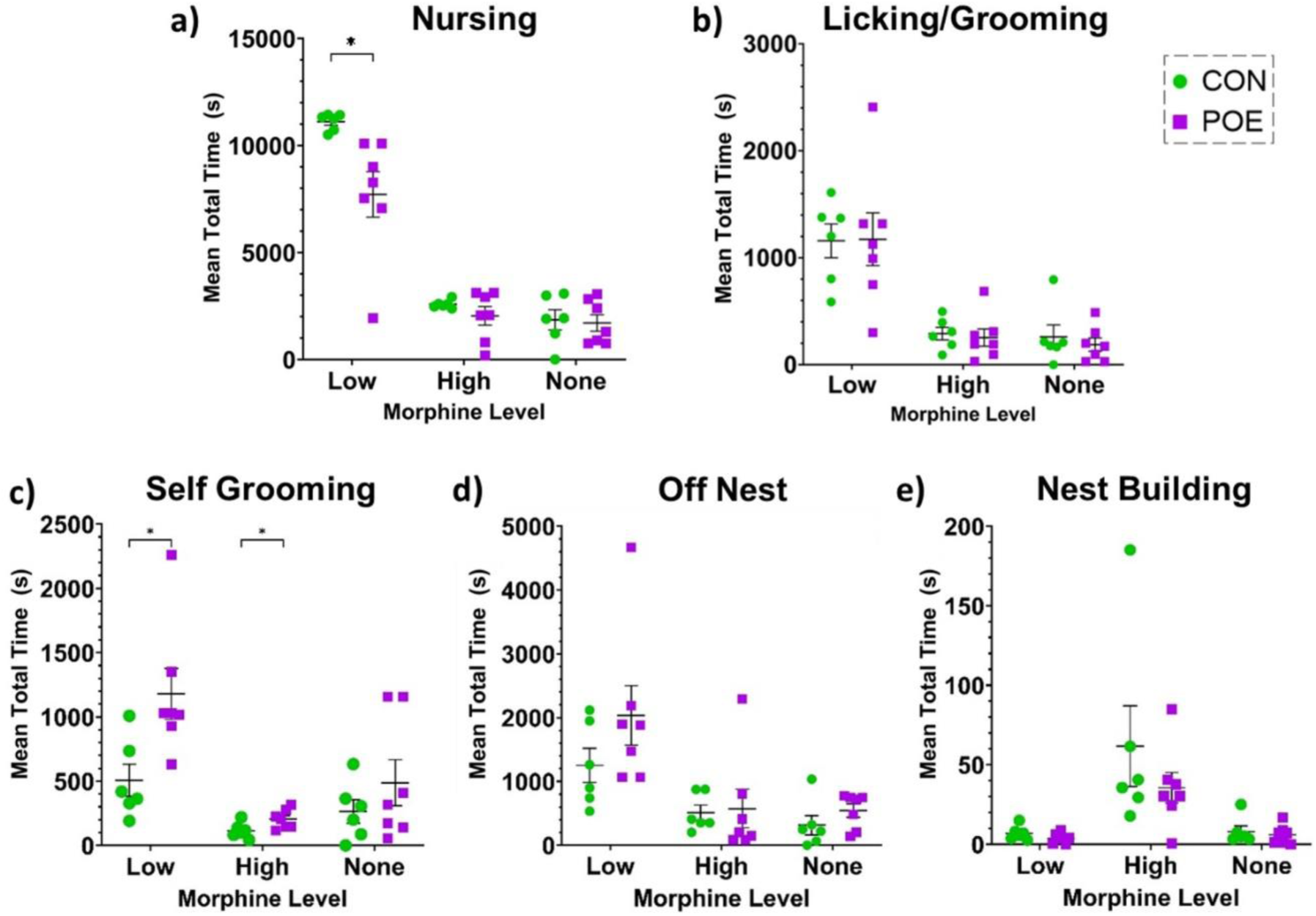
Total time dams spent in each behavioral state was analyzed across morphine levels for nursing. (**a**), licking/grooming of pups (**b**), self-grooming (**c**), off nest (**d**), and nest building (**e**). *Significant difference: Total Time Nursing (CON vs POE low morphine levels), Total Time Self-Grooming (CON vs POE low and high morphine levels). Graphs indicate mean ± SEM.

For self-grooming, there were significant main effects of session and treatment, with POE dams spending more time self-grooming than controls and both groups spending less time self-grooming during afternoon sessions (when morphine levels were high for the POE dams) (F_session_(1.511,16.62)=13.48, p=0.0007; F_treatment_(1,11)=9.759, p=0.0097; Figure 3c). For time off-nest and nest building, there were significant effects of session and no effect of treatment (Off Nest: F_session_(2,22)=19.57, p<0.0001; F_treatment_(1,11)=1.355, p=0.2690; Figure 3d; Nest Building: F_session_(1.038,11.42)=10.56, p=0.0070; F_treatment_(1,11)=1.504, p=0.2456; Figure 3e).

To determine if overall attention to pups is altered by chronic morphine, we categorized measures of total time spent nursing, licking/grooming, and carrying of pups as “pup-directed behavior”. We observed that POE moms spent significantly less time focused on their pups compared to CON (F_treatment_ (1,11)=2.37, p<0.01; Figure 4a). This was particularly evident when morphine levels were high. Overall, across all sessions, the mean total percentage of time directed towards pups was 86.6% for CON versus 65.4% for POE, a 24.5% decrease.

**Figure 4.**
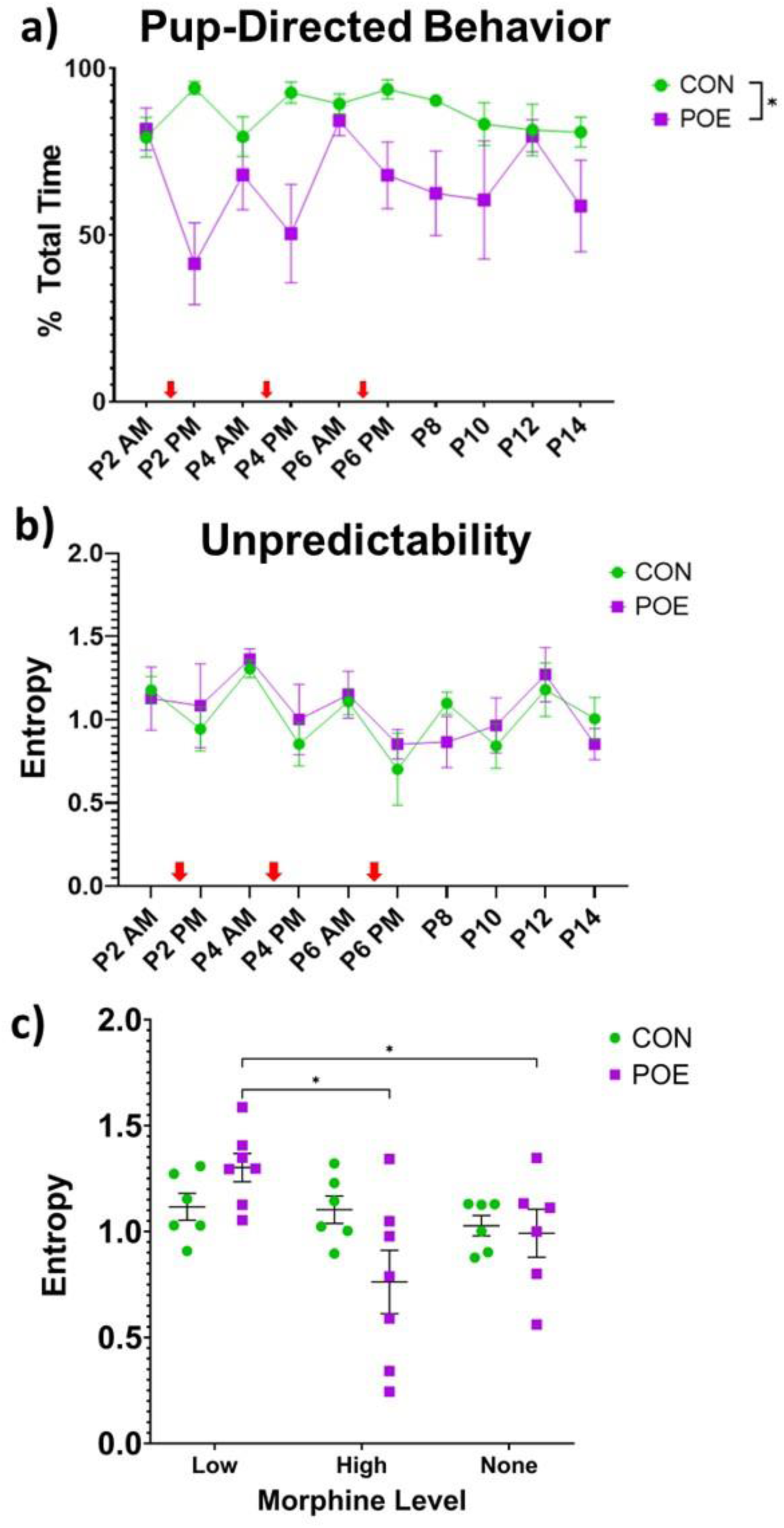
POE dams spent less time performing pup-directed behaviors (nursing, licking/grooming, carrying) compared to CON. (**a**). POE dams also exhibited increased entropy, or unpredictable behavioral patterns, when circulating morphine levels were low, and decreased entropy when morphine levels were high (**b-c**). Red arrows indicate when dams received a bolus of morphine. *Significant difference: %Pup-directed behavior (CON vs POE); Unpredictability (CON vs POE low and high morphine levels). Graphs indicate mean ± SEM.

We next used paired behavioral pattern analysis to calculate an entropy, or unpredictability, score; this score reflects the likelihood of one behavior mode following another. POE dams were less predictable (higher entropy) when morphine was low and more predictable when morphine was high (F_treatment*session_(2,32)=3.96, p=0.03; Figure 4b-c). No significant differences were noted as a function of time for CON dams. Interestingly, POE dams were more unpredictable during morphine withdrawal (Low; AM) than they were following a morphine infusion (High; PM) or after morphine termination (None; P8-P14). This suggests that morphine abstinence/withdrawal in the morning leads to unpredictable behavior, while morphine “intoxication” in the afternoon mutes behavioral variability and increases predictability.

### Sucrose Preference

Our maternal behavior analysis results suggest that POE dams display disrupted maternal care that varies in fragmentation and predictability. Disrupted maternal care has been previously associated with anhedonia or depressive-like behavior in the offspring. Thus, to assess anhedonia in our POE offspring, rats born to either CON or POE dams were tested for their preference for sucrose using two-bottle choice and stress reactivity using the forced swim test.

No effect of sex was observed in sucrose preference, so data were combined. Compared to CON, POE rats showed a decreased preference for sucrose (F_treatment_(1,19)=6.24, p=0.02; Figure 5a; 78.6% CON versus 71.4% POE, 9.2% decrease). POE and CON rats consumed similar amounts of sucrose during habituation, suggesting that preference was unrelated to caloric intake (Figure 5b).

**Figure 5.**
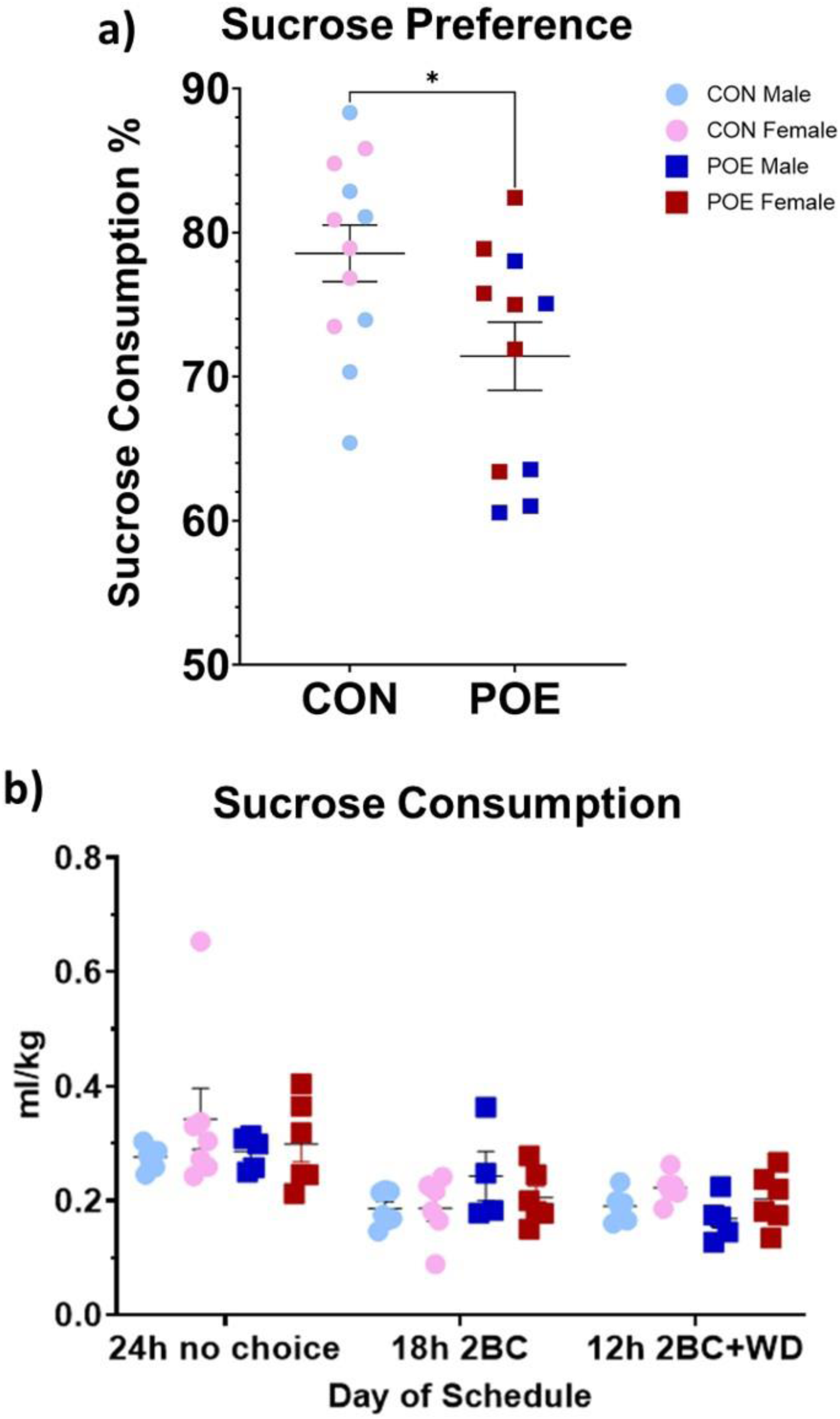
Adolescent offspring were measured for sucrose preference using two-bottle choice for 1% sucrose solution vs water. POE rats exhibited a significant decrease in their sucrose preference compared to CON during testing (**a**). This result was not due to differences in sucrose consumption during reward habituation (**b**). *Significant difference between POE vs CON sucrose preference. Graphs indicate mean ± SEM.

### Acute Stress Reactivity

Approximately two weeks following the sucrose preference test, adult POE and CON rats underwent the forced swim test (FST). We observed no significant effect of treatment on either time spent immobile or latency to immobility (Figure 6a-b). Blood corticosterone levels were determined at baseline, immediately after FST, and 30– and 75-min post-test. There was a significant interaction between treatment and sex (F_treatment*sex_(1,21)=4.73, p=0.04; Figure 6c).

**Figure 6.**
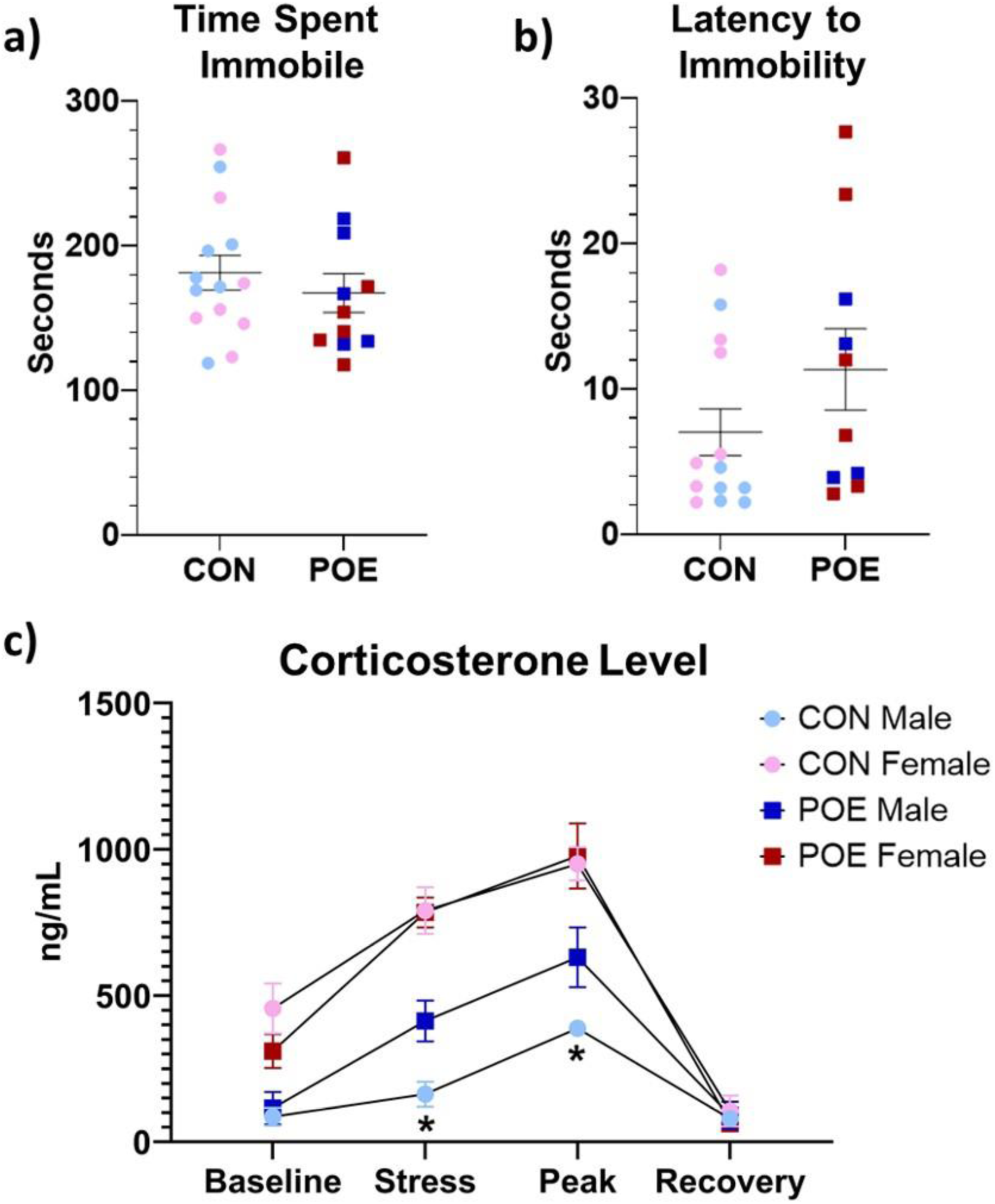
Adult POE and CON performance in the forced swim test (FST). Depression-like behavior, as measured by time spent immobile and latency to immobility, was similar between treatment groups (**a-b**). HPA reactivity, as measured by plasma corticosterone levels across four key timepoints relative to the test, indicated that POE males had an increased stress response to FST, specifically immediately following and at peak (30-min post-test) measures. Female rats exhibited similar corticosterone levels across FST (**c**). *Significant differences between CON vs POE stress corticosterone and CON vs POE peak corticosterone. Graphs indicate mean ± SEM.

POE males exhibited higher plasma corticosterone levels immediately after and 30-min post-FST compared to CON males (F_treatment_(1,28)=10.29, p<0.01). No effect of treatment was observed in females. Overall, FST-induced corticosterone release was higher in females than in males (F_sex_(1,21)=65.44, p<0.01).

### Adult Social Interaction

Our lab has previously reported reduced juvenile play in POE female rats (Harder et al., 2023). Adult POE and CON rats were assessed for time spent socially interacting to determine if social aversion extends later into life. POE rats spent significantly less time interacting with cage mates compared to CON (F_treatment_(1,18)=9.27, p<0.01; 382s CON versus 341s POE, 10.7% decrease; Figure 7a). Females overall spent less time interacting with their cage mates than males (F_sex_(1,18)=11.10, p<0.01; 380s Male versus 335s Female, 11.8% decrease).

**Figure 7.**
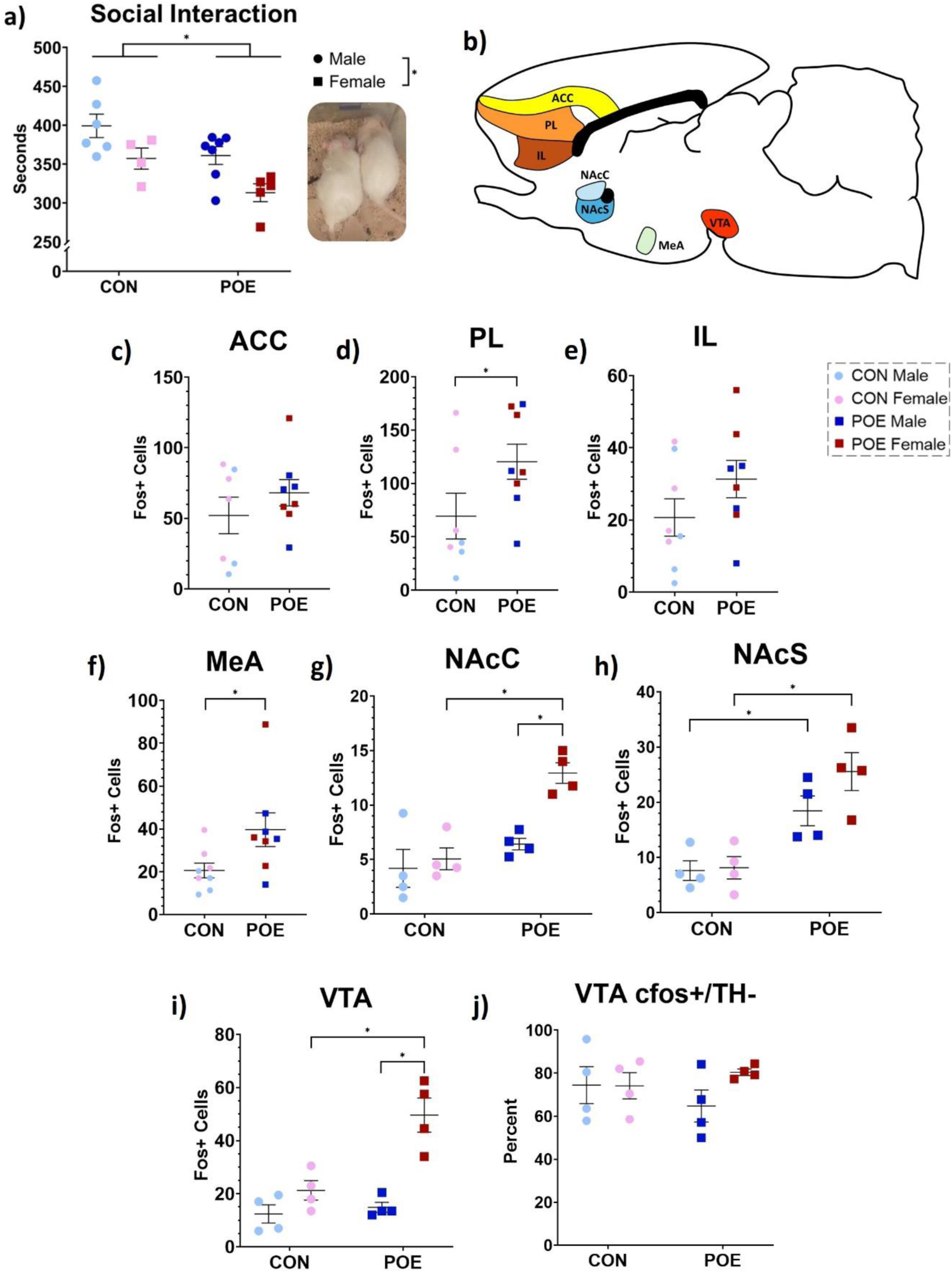
Adult POE and CON rats were tested for social interaction time as a measure of anhedonic behavior. Dual-housed rats were isolated for 24 hours and then reunited with cage mates. POE rats significantly reduce their social interaction time compared to CON rats. Female rats also spent less time with their cage mates overall (**a**). Rats were sacrificed 30-min post-test to determine cellular activity levels in brain regions governing social reward and aversion (**b**). Post-social interaction Fos expression was unchanged in the anterior cingulate cortex (ACC; **c**), increased in the prelimbic cortex (PL; **d**), unchanged in the infralimbic cortex (IL; **e**), and increased in the medial amygdala (MeA; **f**) in POE rats. POE sex-specifically increased Fos^+^ cell counts in nucleus accumbens core and shell, as well as ventral tegmental area (NAcC, NAcS, VTA; **g-i**). Percentage of Fos^+^ cells in the VTA that were nondopaminergic was comparable across groups (**j**). *Significant differences: Social Interaction Time (CON vs POE; males vs females); PL Fos (CON vs POE); MeA Fos (CON vs POE); NAcC Fos (POE females vs all other groups), NAcS Fos (CON males vs POE males; CON females vs POE females), VTA Fos (POE females vs all other groups). Graphs indicate mean ± SEM.

Next, we examined Fos expression in a cohort of rats 90 min after the social interaction test. Positive cell counts were conducted in seven brain regions associated with social reward and aversion (Figure 7b). The number of Fos^+^ cells in an additional four regions averaged less than 10 and were excluded from our analysis: basolateral amygdala, central amygdala, lateral habenula, and medial habenula. Expression of Fos was significantly higher (86% increase) in the prelimbic cortex (PL) of POE rats compared to CON, but not in anterior cingulate (ACC) or infralimbic (IL) cortices (F_treatment_(1,11)=5.164, p=0.04; Figure 7c-e). The number of Fos^+^ cells was also 93% higher in the medial amygdala (MeA; p=0.04; Figure 7f).

Fos expression within the mesolimbic circuit was greater in POE rats than CON rats. Specifically, Fos expression was significantly higher in the NAc core (NAcC), NAc shell (NAcS), and VTA of POE rats compared to CON (NAcC: F_treatment_(1,12)=19.72, p<0.01; NAcS: F_treatment_(1,12)=30.18, p<0.01; VTA: F_treatment_(1,12)=13.57, p<0.01; Figure 7g-i).

Dopaminergic signaling within the VTA is critical for encoding reward, while nondopaminergic signaling, from either GABAergic or glutamatergic interneurons, modulates dopamine activity to dampen reward encoding. Thus, we examined whether the observed Fos^+^ cells in the VTA were co-localized with the dopaminergic marker tyrosine hydroxylase (TH).

Treatment had no significant effect on dopaminergic Fos^+^ cell counts in both POE and CON groups, with the majority of Fos^+^ cells negative for TH (Figure 7j).

## Discussion

The present study aimed to determine the effects of perinatal morphine on maternal care and anhedonia in the offspring. POE dams displayed longer bouts of pup– and self-grooming prior to morphine infusions, which may indicate morphine withdrawal. In the present study, the dosage of morphine was gradually decreased over roughly 10 days surrounding birth to minimize severe spontaneous withdrawal in both dams and pups. Consistent with this, POE dams did not display overt signs of withdrawal, such as wet dog shakes or teeth chattering. At P8, the first full day without morphine, longer bouts of self– and pup-grooming were observed in POE dams. Increased grooming is indicative of withdrawal and is suspected to be a compulsive coping mechanism for abstinence-induced stress or anxiety (Goeldner et al., 2011; Kalueff & Tuohimaa, 2005; Lalanne et al., 2017; Lutz & Kieffer, 2013; Schulteis et al., 1994). POE dams also had longer off-nest bouts at P8. As with increases in grooming at this timepoint, this measure could reflect social avoidance previously reported to accompany rodent opioid withdrawal (Ozdemir et al., 2024). Together, these data suggest that our POE dams did experience withdrawal, which would induce additional stress for both the dam and her offspring.

Overall, our results indicate that gestational opioid exposure induces suboptimal maternal care as POE dams exhibited shorter average bouts of nursing, particularly following morphine administration. These results are consistent with our previous study reporting decreased time spent nursing and increased time licking/grooming pups in POE litters, particularly after the dam receives morphine (Harder et al., 2023). Interestingly, control dams exhibited longer bouts of nursing during P2-P6 afternoon “high morphine” observations than P8-P14 midday observations. This may be explained by decreases in nursing bout length that occurs in the five days after birth; however, hour-to-hour circadian variation in nursing bout length has not been studied (Lapp et al., 2023). Nursing behavior is necessary for offspring, providing both nutritive value and maintenance of body temperature, thus avoiding hypothermia associated with morphine withdrawal (Belknap, 1989; Eriksson & Rönnbäck, 1989). Clinically, both breastfeeding and skin-to-skin contact, even during ongoing opioid maintenance therapy, improves infant health and reduces the need for direct pharmacological treatment in newborns (Abdel-Latif et al., 2006; Atwood et al., 2016; Demirci et al., 2015; Holmes et al., 2016; Kocherlakota, 2014; MacMillan et al., 2018; Patrick et al., 2016; Pritham, 2013; Short et al., 2016; Welle-Strand et al., 2013). Thus, our observation of fragmented maternal care by POE dams, vital for pup survival, is of particular concern for the long-term outcomes of POE offspring.

Compared to CON dams, POE dams varied in the predictability of their maternal behavior, such that morning care (prior to morphine) was less predictable than afternoon care (following morphine). The unpredictability leading up to the morphine infusion may signal reward anticipation or withdrawal; in contrast, the increased predictability of maternal care can be explained by a significant reduction in the different types of behaviors displayed during high levels of morphine. Indeed, during sessions of high morphine levels, POE dams spent significantly less time performing actions that were directed towards pups and more time self-grooming or off-nest. Additionally, the notably broader range in unpredictability scores of the POE dams across all morphine levels (ranging from 0.2 to 1.6 compared to 0.9 to 1.3 for CON) highlights the variability of morphine-exposed dams. This high level of variability would disrupt the pups’ ability to predict maternal care across the first postnatal week and likely hinder reward processing later in life (Novick et al., 2018).

Unpredictable and fragmented maternal care produces postpartum stress that is associated with increased depression, anxiety, and anhedonia in the offspring at both clinical and preclinical levels (Bolton et al., 2018, 2018; Coplan et al., 1996; Levis et al., 2022; Spadoni et al., 2022). In rodents, this is characterized by reduced engagement with rewarding stimuli. For example, suboptimal care induced by resource scarcity reduces socialization and sucrose preference in Sprague Dawley rats (Bolton et al., 2018; Molet et al., 2016). Here, we report adolescent POE rats have reduced preference for sucrose and spend less time socially interacting with cage mates in adulthood. These results and our previous findings of reduced juvenile play suggest POE may promote anhedonia in the offspring(Harder et al., 2023). These behavioral changes likely result from a combination of indirect drug exposure (through the placenta and postnatal milk) and receipt of suboptimal maternal care.

In the present study, POE males had higher levels of CORT following the FST, consistent with previous data reporting increased corticosterone release during stress response to lipopolysaccharide in males but not females following prenatal opiate exposure (Hamilton et al., 2005). Interestingly, despite this sex-specific physiological response to the acute stressor, we found no treatment differences in behavioral response to the FST. Previously in our lab, we have reported a male-specific reduction of corticosterone release following adult FST when a single dose of morphine was given on the day of birth, indicating that chronic versus acute morphine exposure in early life may have opposing effects on stress reactivity (Victoria et al., 2014). Furthermore, the literature regarding the impact of gestational opioid exposure on FST performance seems to highlight the importance of testing age in the consistency of results. For example, Ahmadalipour & Rashidy-Pour (2015) reported prenatal morphine exposure did not impact FST performance at P46, while Chen et al. (2022) reported prenatal morphine exposure in females decreased immobility in the FST at P32 yet increased immobility at P54. Thus, additional FST at multiple timepoints may better elucidate the impact of our POE model on acute stress reactivity across the lifespan.

Gestational morphine resulted in reduced social interaction in adults in the current study, consistent with our previous report of reduced social play behavior in adolescent POE females and reduced time spent together in POE males and females (Harder et al., 2023). In line with our findings, low sociability and social avoidance have been reported in rodents following chronic opioid use in adulthood (Goeldner et al., 2011; Lalanne et al., 2017; Lutz & Kieffer, 2013). However, contrary to our findings, Hol et al. (1996) showed that social play, grooming, and approach behaviors were increased in adolescent and adult rats with prenatal morphine exposure. Notably, the opposing directionality in sociability following early life morphine exposure is likely due to inconsistent dosing timelines, which has contributed to many other experimental outcome discrepancies in the field, particularly regarding reward behavior (Grecco & Atwood, 2020). More specifically, previous studies examining the long-term impact of gestational morphine exposure on social play typically administered daily injections of morphine within a truncated period, typically E8-21, or utilized extended dosing regimens that extended until weaning (Buisman-Pijlman et al., 2009; Hol et al., 1996).

In humans, exposure to opioids *in utero* reduces socialization, coinciding with cognitive deficits and increased anxiety that augment social disorders (Azuine et al., 2019; Larson et al., 2019). Gestational exposure to morphine also reduces social competency scores in infancy and early childhood, increasing the risk of developing ADHD and autism spectrum disorder (Hunt et al., 2008; Sandtorv et al., 2018). Social response in rodents has been altered via pharmacological manipulation of MOR or dopaminergic neurotransmission, highlighting the roles of these systems in social behavior (Beatty & Costello, 1982; Niesink & Van Ree, 1989; Trezza & Vanderschuren, 2008, 2009). Opioidergic and dopaminergic brain regions implicated in social reward show elevated Fos expression following social testing; aberrant functional connectivity in rats that received poor maternal care and early life stress has also been reported (Bolton et al., 2018; Gómez-Gómez et al., 2019; Mejía-Chávez et al., 2021). We found significantly elevated Fos expression in the VTA and NAc in POE rats compared to CON, especially females, the majority of which was nondopaminergic. While aversive stimuli can trigger dopaminergic signaling as an associative learning response, nondopaminergic VTA signaling promotes feelings of anxiety and underlies defensive behavior (Barbano et al., 2020; Cohen et al., 2012; Tan et al., 2012; Zhou et al., 2019). Together with our data, this suggests that POE decreases sociability in a familiar setting during adulthood and increases nondopaminergic activity in the mesolimbic pathway that typically contributes to aversion.

Reduced responses to natural rewards, such as social interaction and sucrose, are a hallmark of anhedonic behavior and are often accompanied by changes in opioidergic or dopaminergic signaling (Der-Avakian & Markou, 2012; Kringelbach & Berridge, 2016; Nestler & Carlezon, 2006). Previous studies have reported activation of the μ-opioid receptor (MOR) in the habenula suppresses depression-like behavior. Specifically, Park et al. (2024) found that DAMGO activation of MOR attenuated aversion and decreased immobility in the forced swim test following acute traumatic stress in rats. Similarly, Bailly et al. (2023) reported that optical stimulation of MOR+ habenular neurons increased both avoidance behavior and immobility in the tail suspension task. These findings, along with our previous report of reduced MOR in the MHb of POE rats, suggest that perinatal morphine reduces MOR-mediated inhibition of habenula neurons, thus facilitating depressive-like behavior (Searles et al., 2023). This lack of opioid inhibition and the currently reported nondopaminergic VTA activation following social interaction identify two independent cellular pathways through which POE-induced alterations in neural signaling promote an anhedonic phenotype.

In summary, we demonstrate that perinatal morphine, which is known to alter preference and motivation for drugs of reward, produces a decreased preference for both sucrose solution and social stimulation. We also show that, in males, POE generates hypothalamic-pituitary-axis hyperactivity during acute stress. These results highlight underlying depression-like symptoms that are produced by the physical and psychological stress of drug exposure in both rodent models and clinical populations. Further, we show that morphine exposure across pregnancy generates fragmentation and inconsistency in maternal care that has been demonstrated in other models of early life adversity to contribute to anhedonia.

While not directly measured, we believe the changes in maternal care induced by morphine also contribute to our previously reported changes in alcohol consumption. Future studies will aim to directly manipulate maternal care through cross-fostering and elucidate the extent to which the observed anhedonic profile of POE offspring is driven by suboptimal maternal care.

Additionally, future analysis correlating offspring measures with the severity of maternal care deficits can further delineate the influence of drug use and rearing conditions on offspring outcomes.

## Acknowledgements

This work was supported by the National Institutes of Health (1RO1DA041529) and the Beckman Scholars Program. The funding sources were not involved in study design, data collection, analysis or interpretation, manuscript writing, or the decision to publish this article. The NIDA Drug Supply Program provided morphine sulfate. Dr. Jessica Bolton assisted with maternal behavior analysis at Georgia State University.

